# Geroprotective interventions preserve trabecular bone during ageing in female mice

**DOI:** 10.64898/2026.06.15.732373

**Authors:** Enrico Dall’Ara, Dharshini Sreenivasan, Sara Oliviero, Maya Boudiffa, Richard A. Miller, Miguel Juarez, Ilaria Bellantuono

## Abstract

Geroprotectors extend lifespan and improve several aspects of healthspan, yet their effects on skeletal ageing remain poorly understood. They hold potential advantages over current bone-targeted osteoporosis therapies, as they may simultaneously improve bone, neuromuscular function, and vision, thereby reducing the risk of falls, the major cause of fractures. Here we examined, for the first time, the long-term effects of rapamycin, acarbose, and 17α-estradiol, administered at lifespan-extending doses on trabecular and cortical bone architecture in male and female UM-HET3 mice measured with micro–computed tomography at 12 and 22 months of age. Bayesian modelling analysis reveals that all interventions produced responses in trabecular bone in females at 22 months. These effects were driven mainly by increases in trabecular number, with little evidence for changes in trabecular thickness. In contrast, treatment effects in males were generally negligible. Cortical responses were modest. Moderate increases in cortical area fraction were observed in females treated with rapamycin or 17α-estradiol at 22 months, whereas cortical thickness remained largely unchanged, suggesting a geometrical rather than anabolic effect. Interestingly, geroprotectors’ strongest skeletal responses in females contrasts with the predominantly male-biased lifespan extension reported for acarbose and 17α-estradiol, suggesting differential mechanisms mediating lifespan extension and bone structure preservation.

## Introduction

Geroprotectors target mechanisms of ageing to delay its detrimental effects (1). They extend lifespan and improve healthspan including physical performance, and age-related eye conditions such as cataract (2). However, their effects on bone properties during ageing remain poorly understood, despite significant age-related bone loss leading to increased bone fragility and fracture risk.

Bone loss affects around one in five men and one in three women over the age of 50 (3). It results from an imbalance between increased bone resorption and a reduced capacity for bone formation with age (4). Current treatments for individuals at high risk of fracture include antiresorptive agents (bisphosphonates or, if contraindicated, denosumab) and anabolic therapies (teriparatide, abaloparatide, and romosozumab) (5). These treatments reduce vertebral fracture risk by 40-70% and non-vertebral fracture risk by 20-40% depending on the agent used (6). However, residual fracture risk reflects not only bone weakness but also age-related decline in muscle function, balance and vision. Interventions that simultaneously target these components would therefore be highly beneficial.

Here we assess the effects of 3 geroprotective interventions, Acarbose (ACA), 17α-Estradiol (17αEST) and rapamycin (Rapa), on bone architecture following prolonged administration at lifespan extending doses. We take advantage of cohorts established at the University of Michigan, as part of the National Institute on Aging Interventions Testing Program(7) ACA is a bacterial-derived α-glucosidase inhibitor that slows the breakdown of complex carbohydrates to glucose thereby reducing post-prandial glucose excursions (8). It extends median and maximum lifespan in UM-HET3 mice with stronger effects in males and improves liver, kidney and neuromuscular function in females (9).

17αEST is a naturally occurring stereoisomer of 17βEST that is non-feminizing, due to reduced activation of classical oestrogen receptors (ERα and Erβ) (10). It extends the median lifespan in male UM-HET3 mice (11) and ameliorates age-related metabolic dysfunction, inflammation (12) and muscle atrophy (13).

Rapamycin is an inhibitor of mammalian target of rapamycin (mTOR) signalling and has been shown to extend lifespan and delay multiple age-related pathologies in both male and female mice (1). Previous studies suggest that rapamycin can attenuate trabecular bone loss under conditions of skeletal challenge, including ovariectomy iron overload, cancer-associated bone disease and ageing (14). However, most studies have focused on trabecular bone with limited assessment of cortical bone, which is also critical for bone strength. Notably, short term treatment has not been associated with improvement in cortical bone (14). Long-term studies assessing both cortical and trabecular compartment are therefore required to determine the full impact of rapamycin on bone ageing.

## Methodology

### Animal experiment

Male and female UM-HET3 mice, which were generated through a cross between (BALB/cBy x C57BL/6)F1 females and (C3H/He x DBA/2)F1 males, were used in this study.

They are the preferred strain used by the National Institute of Aging Interventions Testing Program because they are considered genetically more heterogenous than other inbred strains, better modelling the genetic variation observed in humans. Male and female UM-HET3 mice were randomised to control or treated groups. The treated mice received either ACA 1000 ppm or Rapa (14 ppm) in their diet starting at 4 months.17αEST was administered starting at 10 months at 14.4ppm in their diet. Animals were sacrificed at 12- or 22-months and the tibias excised for analysis.

### MicroCT imaging and image processing

The mouse tibia was dissected from the euthanized mice and processed for assessing its morphometric properties. The right tibia of each mouse was scanned (VivaCT 80, Scanco Medical, Bruettisellen, Switzerland) *ex vivo* with the following scanning protocol (55 kVp, 145 μA, 10.4 μm voxel size, 100 ms integration time, 32 mm field of view, 750 projections/180°, no frame averaging, 0.5 mm Al filter) (15). All images were reconstructed using the software provided by the manufacturer (Scanco Medical AG) and applying a polynomial beam hardening correction based on a phantom of 1200 mg HA/cc density, which has been shown to improve the local tissue mineralization measurement (16). MicroCT images were used to estimate the following parameters of interest: morphometric parameters of the proximal metaphyseal trabecular bone and of the diaphyseal cortical bone (15). For trabecular morphological measurements, a volume of interest (VOI) of 1 mm was selected below the growth plate with an offset of 0.2 mm from a reference slice, identified as the point where the medial and lateral sides of the growth plate merged (17). Trabecular bone was separated from the cortical shell by manual contouring every five slices in each microCT stack of 2D images. Linear interpolation was applied between slices. A single level threshold, calculated as the average of the grey levels corresponding to the bone and background peaks in the image histogram (18) was used to segment the images. A despeckling filter was applied to remove 3D bone volumes smaller than 10 voxels. Trabecular bone volume fraction (Tb.BV/TV), thickness (Tb.Th.), separation (Tb.Sp.) and number (Tb.N.) were computed for each trabecular bone region of interest (ROI).

Cortical morphological measurements were performed on a ROI of 1 mm, centred at the tibial midshaft [Oliviero2022]. After segmentation, pores within the cortex were removed by applying a closing function (2D round kernel, radius equal to 10 pixels).

All morphometric analyses were performed using CTAn (v1.18.4.0, Skyscan-Bruker, Kontich, Belgium).

### Statistical analysis

Statistical analyses were performed to assess the effects of pharmacological treatment on trabecular and cortical bone parameters as a function of sex and age. Data exploration revealed heterogeneity of variance across experimental groups, precluding the use of standard parametric approaches assuming homoscedasticity. Therefore, a Bayesian modelling framework was adopted to allow group-specific variance and to provide direct probabilistic estimates of treatment effects in small groups (19, 20).

Bayesian models restricted to untreated control animals were used to assess age-related changes in bone parameters relative to the 4-months baseline. This analysis estimated posterior mean differences between baseline and subsequent time points (12 and 22 months) as a function of sex.

For each bone outcome, treatment effects were estimated relative to control animals, stratified by sex (female, male) and age (12 and 22 months). Trabecular and cortical parameters were analysed in separate models.

Trabecular number and thickness were log-transformed prior to analysis to improve model fit, while cortical relative density was analysed on the odds (logit) scale to account for the bounded nature of the variable. All other outcomes were analysed on their original scale.

For baseline ageing analysis, posterior mean differences relative to 4-month reference group were estimated in the same manner.

Model results are reported as posterior mean differences between treatment and control groups, together with 95% credible intervals, defined as the range containing 95% of the posterior probability distribution for the estimated effect. Credible intervals were used to describe the uncertainty of estimated treatment effects.

To further quantify the strength and consistency of treatment effects, standardized effect sizes were calculated and evaluated using regions of practical equivalence (ROPE). Effect sizes were classified as negligible (absolute value < 0.2), small (0.2–0.5), medium (0.5–0.8), or large (> 0.8). Posterior probabilities associated with each effect-size category were used to assess the likelihood that observed differences represented biologically meaningful effects.

All results are presented as posterior estimates with associated uncertainty. No null-hypothesis significance testing or p-values were used.

## Results

### UM-HET3 mice undergo bone loss with age

Trabecular bone volume fraction (BV/TV) declined substantially with age in both female and male mice. In females, posterior estimates indicated reductions of 6.69% at 12 months relative to baseline (95% credible interval: −7.94 to −5.39) and 8.06% at 22 months relative to baseline (95% credible interval: −9.13 to −7.05) (Fig1). These findings indicate that the majority of trabecular bone loss in females occurred before 12 months of age. In males, trabecular bone loss at 12 months was smaller than in females, with the sex × time interaction indicating a difference of 1.70% (95% credible interval: 0.07 to 3.37). Estimated trabecular bone loss in males was therefore 4.99% at 12 months and 7.50% at 22 months relative to baseline, indicating a more gradual decline with age. By 22 months, the difference between sexes was reduced and associated with greater uncertainty (sex × month22 interaction: 0.56, 95% credible interval: −0.96 to 2.07).

Cortical Area/Tissue area declined substantially with age in both female and male mice. Bayesian modelling identified reductions of 6.14% at 12 months relative to baseline (95% credible interval: −7.89 to −4.32) and 10.06% at 22 months relative to baseline (95% credible interval: −11.76 to −8.35) (Fig. 2). No evidence for sex-dependent differences in the trajectory of cortical bone loss was identified, although male mice exhibited overall lower cortical bone values than females (mean difference −3.72, 95% credible interval: −5.27 to −2.33). Cortical bone loss continued after 12 months, with an additional decline of approximately 3.92% occurring between 12 and 22 months.

**Fig. 1.**
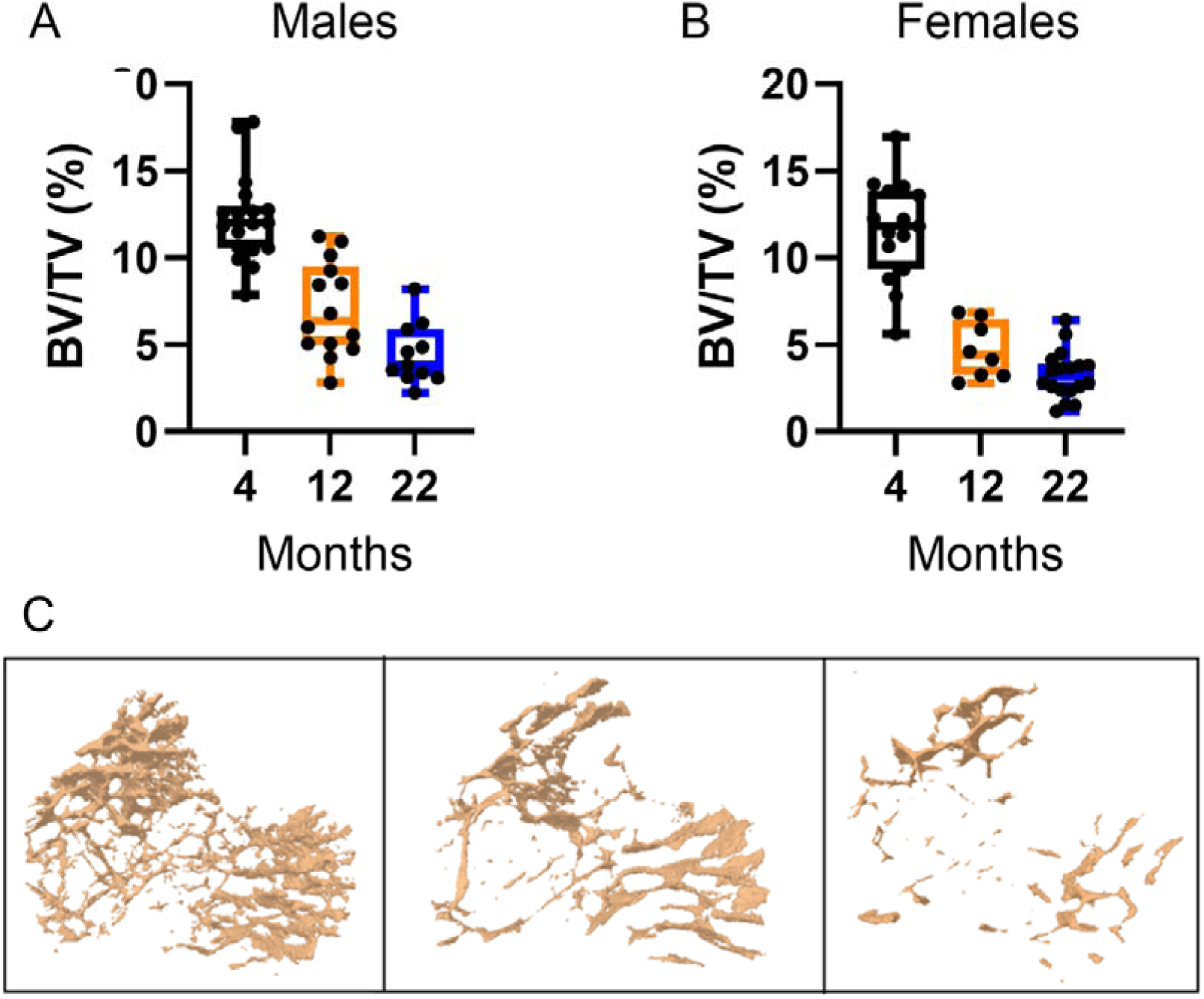
UM-HET3 mice undergo trabecular bone loss with age. Box plot shows the median (center line), interquartile range (box) and maximum and minimum value (whiskers). Each dot represents one animal. A) Trabecular Bone Volume fraction (BV/TV) in male UM-HET3 mice at 4, 12 and 22 months of age; B) Trabecular Bone Volume fraction (BV/TV) in female UM-HET3 mice at 4, 12 and 22 months of age; C) A representative 3D micro-CT reconstruction of trabecular bone from the tibia of a UM-HET3 male mouse at 4 (left panel), 12 (middle panel) and 22 months (right panel).

**Fig. 2.**
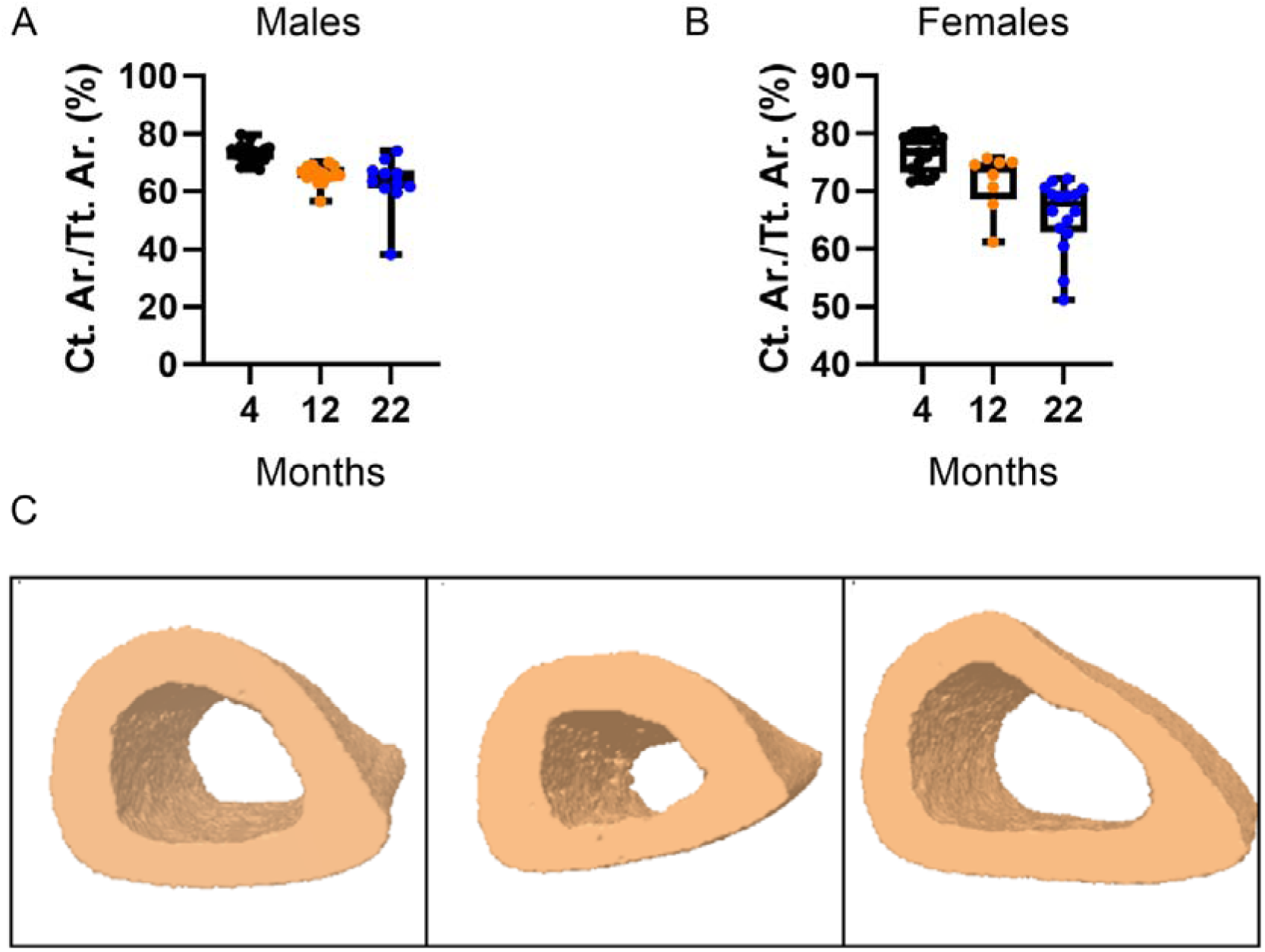
UM-HET3 mice undergo cortical bone loss with age. Box plot shows the median (center line), interquartile range (box) and maximum and minimum value (whiskers). Each dot represents one animal. A) Cortical area fraction (Ct Ar/T Ar) in male UM-HET3 mice at 4, 12 and 22 months of age; B) Cortical area fraction (Ct Ar/T Ar) in female UM-HET3 mice at 4, 12 and 22 months of age; C) A representative 3D X-ray micro-CT reconstruction of cortical bone from the tibia of a UM-HET3 male mouse at 4 (left panel), 12 (middle panel) and 22 months (right panel).

### Interventions increase trabecular bone volume in female mice

A schematic representation of the experimental design is in Fig.3. Trabecular bone volume fraction (BV/TV) exhibited sex- and age-dependent responses to interventions, with statistically supported effects observed predominantly in female mice (Figure 4A-B; Tables 1–2).

**Fig. 3.**
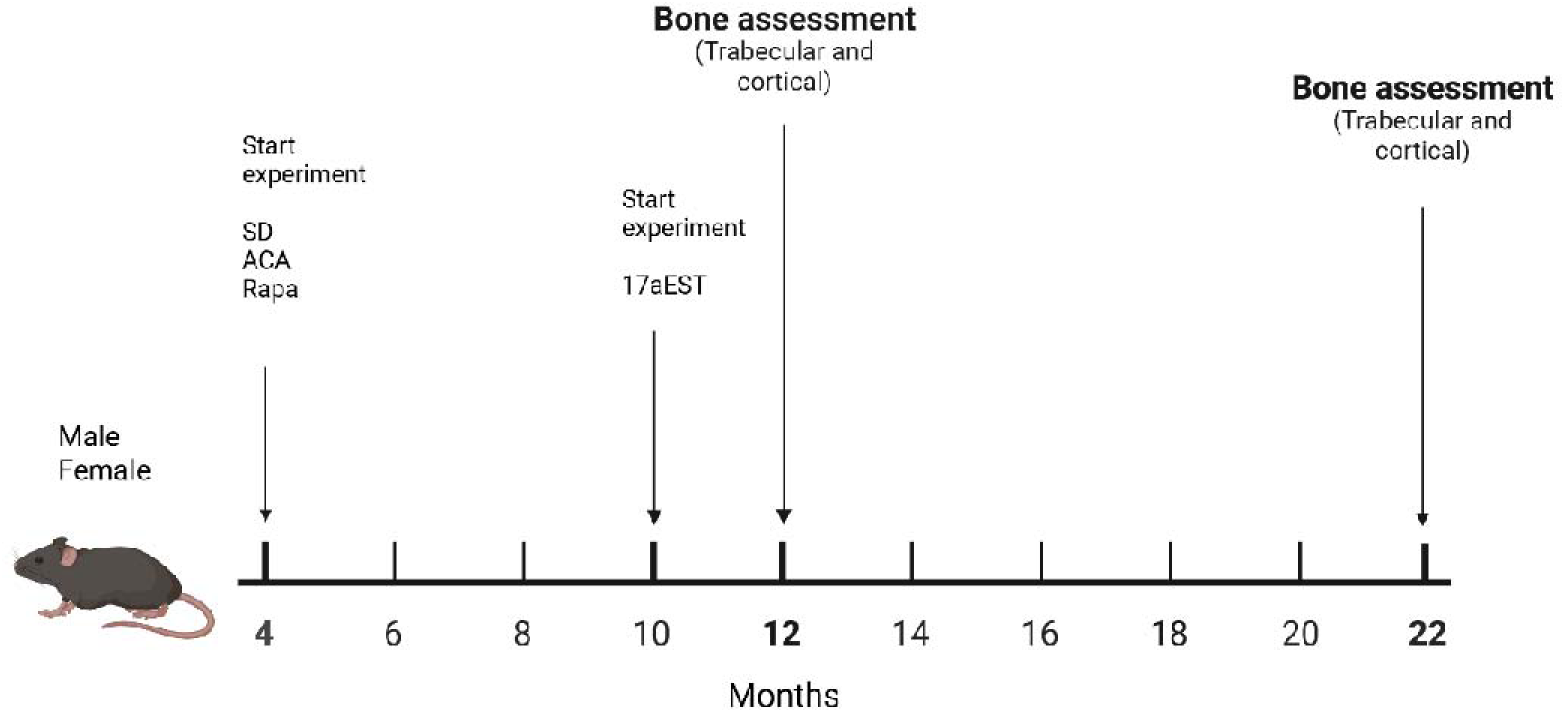
Graphic representation of the experimental n Timelines. This was a cross sectional study comprising one group of mice culled at 4 months, 4 groups of mice (one group/intervention plus one on standard diet used as control) culled at 12 months and 4 more groups culled (3 interventions plus control) at 22 months

**Fig. 4.**
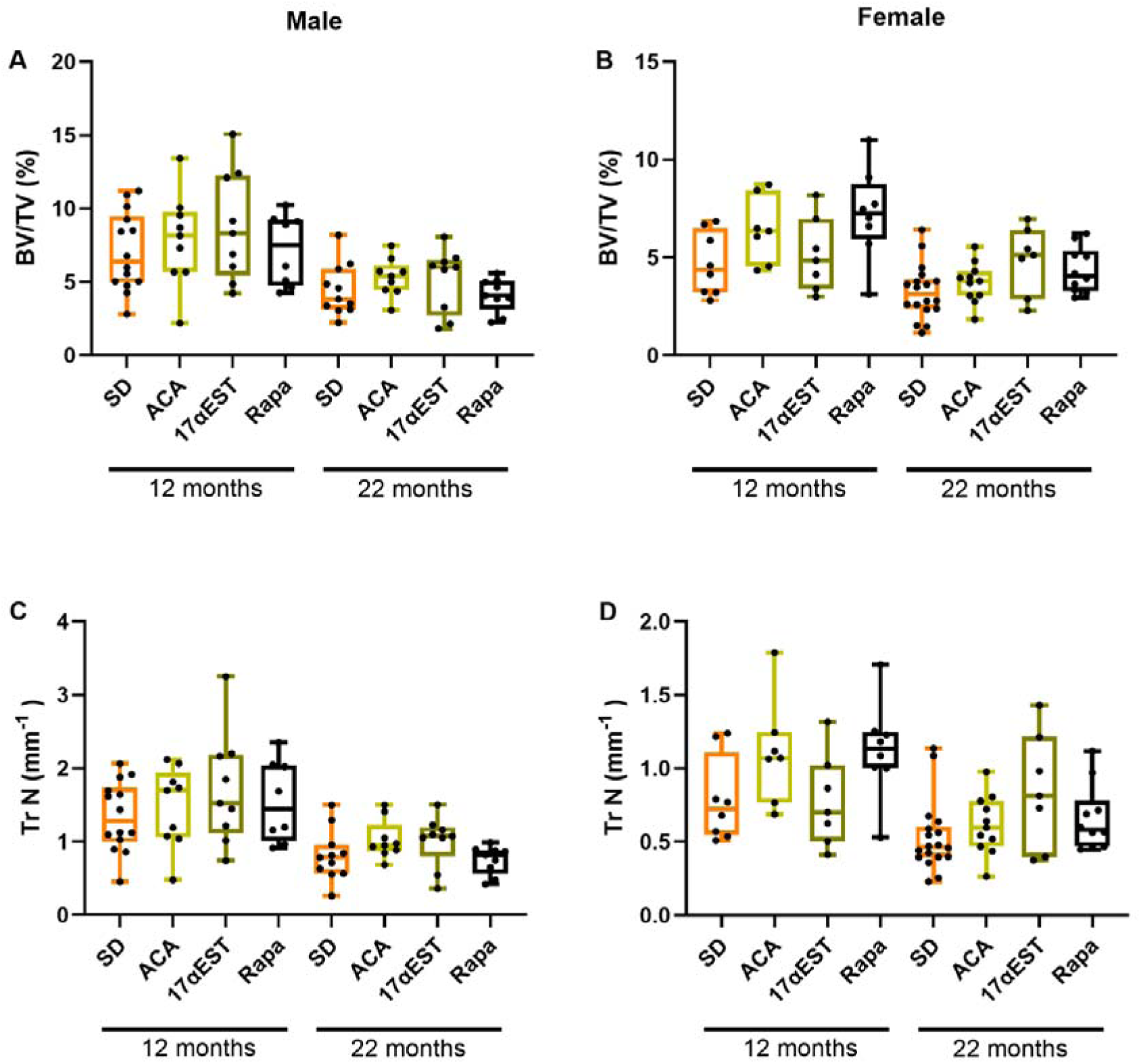
Geroprotective interventions increase trabecular bone volume fraction and trabecular number in female UM-HET3 mice. Box plot shows the median (center line), interquartile range (box) and maximum and minimum value (whiskers). Each dot represents one animal. A) Bone Volume fraction (BV/TV) in male UM-HET3 mice receiving standard diet (SD), Acarbose (ACA), 17α-estradiol (17αEST), rapamycin (Rapa), analysed at 12 or 22 months of age. B) Trabecular BV/TV in female UM-HET3 mice exposed to the same interventions; C) Trabecular Number (Tr N) in the same male mice shown in panel A. D) Tr N in the same female mice shown in panel B.

**Table 1.** Posterior probabilities of standardised effect sizes for trabecular bone volume fraction (BV/TV) relative to standard diet (SD), classified according to regions of practical equivalence (ROPE) by sex and age. Rope categories correspond to standardised effects sizes: negligible (absolute value < 0.2), small (0.2–0.5), medium (0.5–0.8), or large (> 0.8). The category with the highest probability in each group is highlighted. 17aEST, 17-α-estradiol; ACA, acarbose; Rapa, rapamycin

| Age | Intervention | Sex | Large | Medium | Small | Negligible |
| --- | --- | --- | --- | --- | --- | --- |
| 12 | 17aEST | Female | 0.17 | 0.244 | 0.348 | 0.238 |
| 12 | 17aEST | Male | 0.639 | 0.147 | 0.131 | 0.083 |
| 12 | ACA | Female | 0.9 | 0.063 | 0.026 | 0.010 |
| 12 | ACA | Male | 0.372 | 0.213 | 0.246 | 0.169 |
| 12 | Rapa | Female | 0.956 | 0.025 | 0.013 | 0.006 |
| 12 | Rapa | Male | 0.17 | 0.204 | 0.342 | 0.284 |
| 22 | 17aEST | Female | 0.927 | 0.051 | 0.016 | 0.005 |
| 22 | 17aEST | Male | 0.211 | 0.306 | 0.317 | 0.166 |
| 22 | ACA | Female | 0.002 | 0.247 | 0.696 | 0.055 |
| 22 | ACA | Male | 0.361 | 0.429 | 0.174 | 0.036 |
| 22 | Rapa | Female | 0.857 | 0.134 | 0.008 | 0.000 |
| 22 | Rapa | Male | 0.022 | 0.167 | 0.496 | 0.315 |

**Table 2.** Posterior probabilities of standardised effect sizes for trabecular number relative to standard diet (SD), classified according to regions of practical equivalence (ROPE) by sex and age. Rope categories correspond to standardised effects sizes: negligible (absolute value < 0.2), small (0.2–0.5), medium (0.5–0.8), or large (> 0.8). The category with the highest probability in each group is highlighted. 17aEST, 17-α-estradiol; ACA, acarbose; Rapa, rapamycin

| Age | Intervention | Sex | Large | Medium | Small | Negligible |
| --- | --- | --- | --- | --- | --- | --- |
| 12 | 17aEST | Female | 0.000 | 0 | 0.001 | 0.999 |
| 12 | 17aEST | Male | 0.000 | 0 | 0.986 | 0.014 |
| 12 | ACA | Female | 0.000 | 0.751 | 0.248 | 0 |
| 12 | ACA | Male | 0.000 | 0 | 0.007 | 0.993 |
| 12 | Rapa | Female | 0.000 | 0.821 | 0.179 | 0 |
| 12 | Rapa | Male | 0.000 | 0 | 0.755 | 0.245 |
| 22 | 17aEST | Female | 0.001 | 0.883 | 0.116 | 0 |
| 22 | 17aEST | Male | 0.000 | 0 | 0.974 | 0.026 |
| 22 | ACA | Female | 0.000 | 0 | 0.981 | 0.019 |
| 22 | ACA | Male | 0.000 | 0.257 | 0.743 | 0 |
| 22 | Rapa | Female | 0.000 | 0.003 | 0.997 | 0 |
| 22 | Rapa | Male | 0.000 | 0 | 0 | 1 |

At 12 months of age, rapamycin (Rapa) produced a statistically robust increase in trabecular BV/TV in females. The posterior distribution of the standardized effect size relative to standard diet showed a 95.7% probability of a large effect, with an estimated mean increase in BV/TV of 2.46% (95% credible interval [CI]: 0.86 to 4.02). Acarbose (ACA) also increased BV/TV in females at this age, with a 90.0% probability of a large effect and a mean increase of 1.71% (95% CI: 0.52 to 2.87). In contrast, 17α-estradiol (17αEST) showed no statistically meaningful effect in 12-month-old females, with most posterior mass falling within the region of small or negligible effects (probability of a large effect 16.8%). However, the administration started later than the other geroprotectors.

At 22 months of age, in females, Rapa continued to show strong evidence of an increase in trabecular BV/TV, with an 85.6% probability of a large effect and a mean increase of 1.08% (95% CI: 0.72 to 1.42). At this later age, 17αEST also demonstrated a statistically supported effect in females, with a 92.7% probability of a large effect and an estimated increase of 1.51% (95% CI: 0.66 to 2.29). In contrast, ACA showed only weak statistical support at 22 months in females, with the posterior distribution dominated by small effect sizes (probability of a small effect 69.6%).

In male mice at 12 months, none of the interventions produced a statistically convincing increase in trabecular BV/TV. Although 17αEST showed a 63.5% probability of a large effect, the corresponding absolute difference and large 95% confidence interval suggests uncertainty (mean 1.69; 95% CI: −2.41 to 5.77). ACA and Rapa in males were dominated by small or negligible effect size probabilities, with mean differences close to zero.

In males at 22 months, the effects of the interventions on trabecular BV/TV were minimal. For 17αEST and ACA, posterior effect size distributions were centred on small effects, with probabilities for large effects of 20.9% and 36.5%, respectively, and mean BV/TV differences below 1.0% (from 4.78% to 5.13% for 17αEST and remaining 4.78% for ACA). Rapa showed no evidence of a positive effect in males at this age, with a 49.3% probability of a small effect and a mean difference of −0.34% (95% CI: −0.97 to 0.30), indicating compatibility with no change relative to standard diet.

To understand the structural basis of the observed changes in trabecular BV/TV, we next examined trabecular number, thickness and spacing. The observed increases in trabecular BV/TV primarily reflected increases in trabecular number (Fig. 4C-D and table 2) rather than changes in trabecular thickness (Supplementary Fig 2 A-B and supplementary table 1). The changes in trabecular separation were also small or negligible, probably reflecting the overall small and variable number of trabeculae present in the analysed region of interest (Supplementary Fig 1C-D and supplementary table 2).

Taken together, the statistical evidence supports a model in which increases in trabecular bone density are largely restricted to female mice and are driven mainly by an increase in trabecular number, with Rapa exerting consistent effects at both ages, ACA acting primarily at 12 months, and 17αEST showing a delayed effect that becomes apparent only in older females.

### Interventions have modest effects on cortical bone in male and female mice

Cortical bone analysis showed more modest and variable responses to interventions than trabecular bone, with effect sizes generally falling in the small to medium range (Figure 5; Table 3-4).

**Fig. 5.**
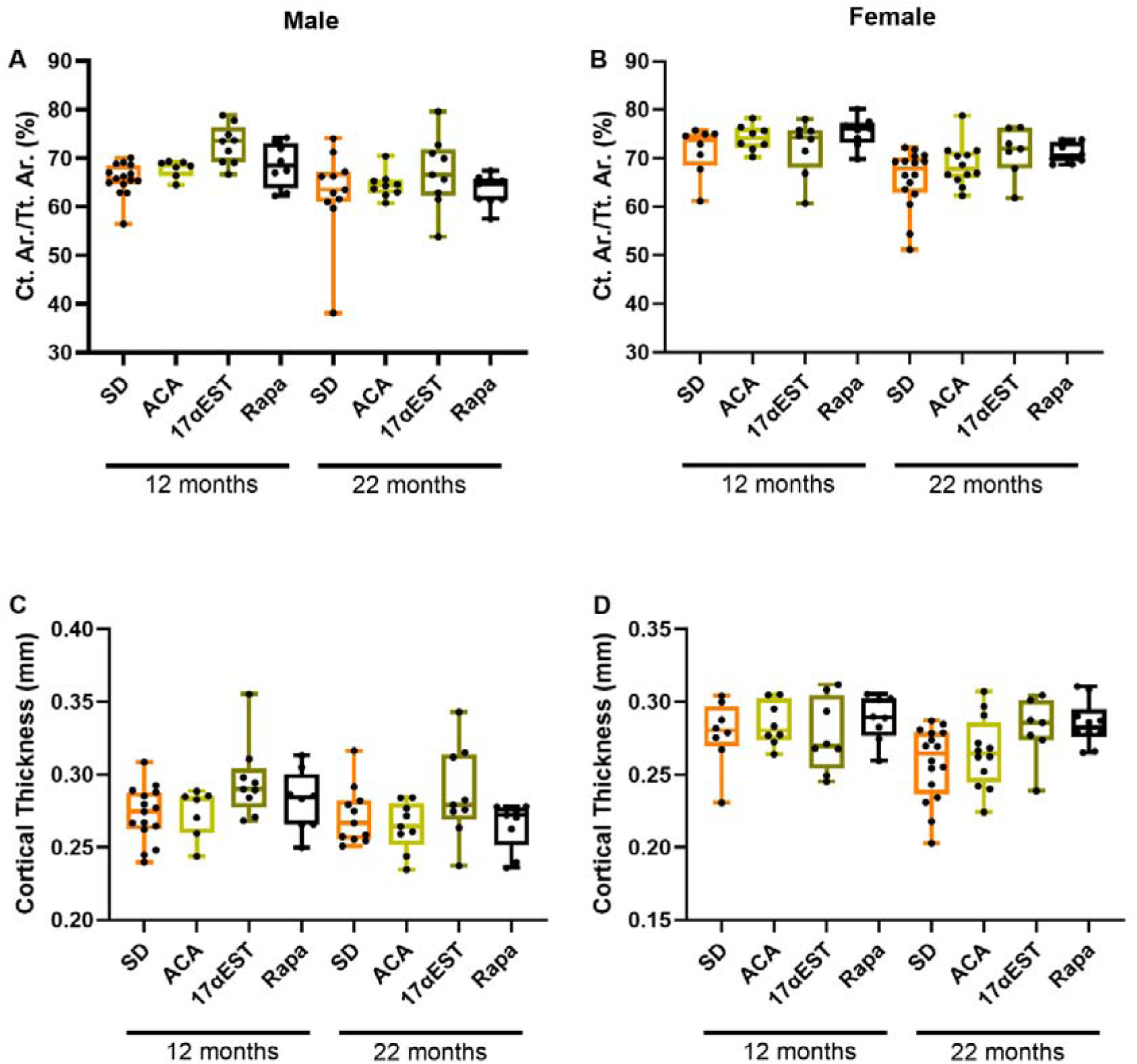
Geroprotective interventions have modest effects on cortical bone in male and female UM-HET3 mice. Box plot shows the median (center line), interquartile range (box) and maximum and minimum value (whiskers). Each dot represents one animal. A) Cortical area fraction (Ct Ar/T Ar) in male UM-HET3 mice receiving standard diet (SD), Acarbose (ACA), 17α-estradiol (17αEST), rapamycin (Rapa), analysed at 12 or 22 months of age. B) Ct Ar/T Ar in female UM-HET3 mice exposed to the same interventions; C) Cortical Thickness (Ct Th) in the same male mice shown in panel A. D) Ct Th in the same female mice shown in panel B.

**Table 3.** Posterior probabilities of standardised effect sizes for cortical area fraction (Cortical Area over Tissue Area) relative to standard diet (SD), classified according to regions of practical equivalence (ROPE) by sex and age. Rope categories correspond to standardised effects sizes: negligible (absolute value < 0.2), small (0.2–0.5), medium (0.5–0.8), or large (> 0.8). The category with the highest probability in each group is highlighted. 17aEST, 17-α-estradiol; ACA, acarbose; Rapa, rapamycin

| Age | Intervention | Sex | Large | Medium | Small | Negligible |
| --- | --- | --- | --- | --- | --- | --- |
| 12 | 17aEST | Female | 0 | 0 | 0.143 | 0.857 |
| 12 | 17aEST | Male | 0.995 | 0.005 | 0 | 0 |
| 12 | ACA | Female | 0 | 0.122 | 0.87 | 0.008 |
| 12 | ACA | Male | 0 | 0 | 1 | 0 |
| 12 | Rapa | Female | 0.086 | 0.865 | 0.049 | 0 |
| 12 | Rapa | Male | 0 | 0.69 | 0.31 | 0 |
| 22 | 17aEST | Female | 0.335 | 0.653 | 0.011 | 0 |
| 22 | 17aEST | Male | 0 | 0.17 | 0.81 | 0.019 |
| 22 | ACA | Female | 0 | 0 | 0.997 | 0.002 |
| 22 | ACA | Male | 0 | 0 | 0.001 | 0.999 |
| 22 | Rapa | Female | 0.258 | 0.742 | 0 | 0 |
| 22 | Rapa | Male | 0 | 0 | 0.014 | 0.986 |

For cortical area fraction (Fig. 5A-B table 3), at 12 months of age, the strongest statistical signal was observed in male mice treated with 17αEST, for which the posterior distribution indicated a 99.5% probability of a large effect size and an approximately 7% increase relative to controls. Rapamycin (Rapa) in males showed a moderate effect (68.6% probability of a medium effect), and an approximate 3% increase in cortical area fraction relative to controls, whereas acarbose (ACA) was associated primarily with small effects. In females at the same age, cortical responses were weaker: Rapa showed a moderate effect similar to that observed in males, ACA was associated with small effects, and 17αEST showed little evidence of a detectable effect.

At 22 months of age, cortical responses were largely restricted to females. In this group, both 17αEST and Rapa showed moderate effect sizes (65.3% and 74.3% probability of medium effects, respectively and approximate increases of ∼5% in cortical area fraction relative to controls), whereas ACA remained confined to small effects. In males at 22 months, posterior effect size distributions were dominated by small or negligible effects across all interventions, indicating little evidence of substantial changes in cortical bone.

The absence of consistent changes in cortical thickness (Fig. 5C–D; Table 4) indicates that these differences in cortical area fraction are likely to reflect limited alterations in cortical geometry rather than increased cortical apposition.

**Table 4.** Posterior probabilities of standardised effect sizes for cortical thickness relative to standard diet (SD), classified according to regions of practical equivalence (ROPE) by sex and age. Rope categories correspond to standardised effects sizes: negligible (absolute value < 0.2), small (0.2–0.5), medium (0.5–0.8), or large (> 0.8). The category with the highest probability in each group is highlighted. 17aEST, 17-α-estradiol; ACA, acarbose; Rapa, rapamycin

| Age | Intervention | Sex | Large | Medium | Small | Negligible |
| --- | --- | --- | --- | --- | --- | --- |
| 12 | 17aEST | Female | 0 | 0 | 0 | 1 |
| 12 | 17aEST | Male | 0 | 0 | 0.387 | 0.613 |
| 12 | ACA | Female | 0 | 0 | 0 | 1 |
| 12 | ACA | Male | 0 | 0 | 0 | 1 |
| 12 | Rapa | Female | 0 | 0 | 0 | 1 |
| 12 | Rapa | Male | 0 | 0 | 0 | 1 |
| 22 | 17aEST | Female | 0 | 0 | 0.543 | 0.457 |
| 22 | 17aEST | Male | 0 | 0 | 0 | 1 |
| 22 | ACA | Female | 0 | 0 | 0 | 1 |
| 22 | ACA | Male | 0 | 0 | 0 | 1 |
| 22 | Rapa | Female | 0 | 0 | 0.991 | 0.009 |
| 22 | Rapa | Male | 0 | 0 | 0 | 1 |

## Discussion

This study shows a consistent effect of these 3 geroprotectors on trabecular bone in UM-HET3 female mice at 22 months of age. The size of the effect differs depending on the intervention. 17αEST and rapamycin showed the greatest increase in trabecular bone density (47% and 34% increase compared to controls of the same age, respectively). The weakest increase at 22 months was observed with Acarbose at 15%. We saw no consistent effect on cortical bone including rapamycin despite the long-term nature of the intervention. Data shows an increase in cortical area fraction but not in cortical thickness in female mice at 22 months suggesting changes in bone geometry rather than in cortical apposition.

The size of the increase observed in trabecular bone with 17αEST is in a similar range than the one reported with senolytics in aged mice (21) and in the lower range of what detected with the administration of Parathyroid Hormone, PTH(1–34), for 4 weeks in 26 months old mice in the study reported by Jilka at al. (2010) (22). PTH(1–34) is a bone anabolic drug which is given a second line therapy in patients affected by osteoporosis. However, in both published studies the analysis was conducted in vertebrae as opposed to our study which analysed changes in tibia. Any comparison should be interpreted cautiously because different skeletal sites produce different responses to mechanical and biochemical stimuli (23, 24).

Interestingly, ACA and 17αEST exhibited different temporal patterns of trabecular bone preservation. ACA showed detectable effects at 12 months, whereas the effects of 17αEST were more evident at 22 months. One possible explanation is the difference in treatment initiation, as ACA administration began at 4 months of age whereas 17αEST treatment commenced at 10 months. The later initiation of 17αEST was intended to avoid potential interference with early hormonal and skeletal development. Given that our ageing analysis indicated that the majority of trabecular bone loss occurred before 12 months, initiation of 17αEST at 10 months may nevertheless have limited its ability to influence the early phase of bone loss. This may be particularly relevant if these interventions act primarily by preserving existing bone architecture rather than through rapid anabolic mechanisms capable of restoring already lost trabecular structure.

Indeed, the structural pattern observed of maintenance of trabecular number, with comparatively little evidence for increased trabecular thickness or cortical apposition is more consistent with preservation of the existing trabecular architecture than with robust anabolic formation. Such effects could plausibly arise through mechanisms including suppression of inflammatory signalling and modulation of osteoclast activity, or preservation of osteoblast progenitor populations. Notably, rapamycin, ACA, and 17αEST have all been reported to suppress inflammatory pathways, including IL-6 and IL-1 signalling, which are associated with enhanced osteoclast-mediated bone resorption (25–28). Previous studies examining short-term administration of Rapamycin in bone in mice suggested that its skeletal effects arise primarily through inhibition of osteoclast formation (14).

Interestingly sex-specific skeletal responses to these interventions did not mirror their effects on lifespan. This divergence is not unprecedented. 17α-estradiol shows male-specific benefits for lifespan (11). It also ameliorates age-associated sarcopenia and improves late-life physical function in males, but not in females or castrated males (13), pointing to dependence on testicular hormones for several male-specific benefits.

In contrast acarbose improves some age-sensitive functional traits in both sexes despite a stronger survival benefit in males. In Herrera et al. (29), acarbose improved several physical-function measures in both sexes, including rotarod performance and grip strength, while some cardiac aging phenotypes were male-specific.

Rapamycin extends lifespan in both sexes, although median lifespan extension is more pronounced in female mice (30, 31). Most studies on the effect of rapamycin on organ function are in male mice (32, 33). However, benefit on organ function is seen in both sexes when tested (32, 33) including muscle function assessed by rotarod and stride length (33).

Our findings therefore suggest that the pathways through which these interventions modulate skeletal aging are at least partly distinct from those underlying their effects on survival. Lifespan extension in laboratory mice often reflects delayed cancer mortality rather than uniform slowing of aging across tissues. Because neoplasia is the predominant cause of death in UM-HET3 mice, sex-specific survival benefits reported for several of these interventions may primarily reflect differences in tumour biology. In contrast, the skeletal responses observed here likely reflect direct effects on bone remodelling and may therefore exhibit a different sex dependence.

Although the skeletal effects observed here were generally more modest than those reported for established bone anabolic therapies, this does not necessarily limit their potential clinical value. Unlike conventional osteoporosis drugs, geroprotective interventions may simultaneously target multiple ageing-related contributors to fracture risk, including neuromuscular decline, and visual impairment, which are important causes of falls. The predominance of skeletal responses in female mice may also be clinically relevant, given the higher prevalence of osteoporosis in post-menopausal women (34). However, the marked sex-specific differences observed between interventions highlight the importance of carefully evaluating tissue- and sex-dependent responses when considering geroprotective strategies for musculoskeletal ageing. Future studies should therefore aim to identify interventions capable of conferring coordinated benefits across multiple organ systems within the same sex.

## Acknowledgments

This grant was funded by BBSRC grant N BB/R001510/1 and by NIH grant AG022303.

