## Supplemental Fig 1 for "Geroprotective interventions preserve trabecular bone during ageing in female mice"

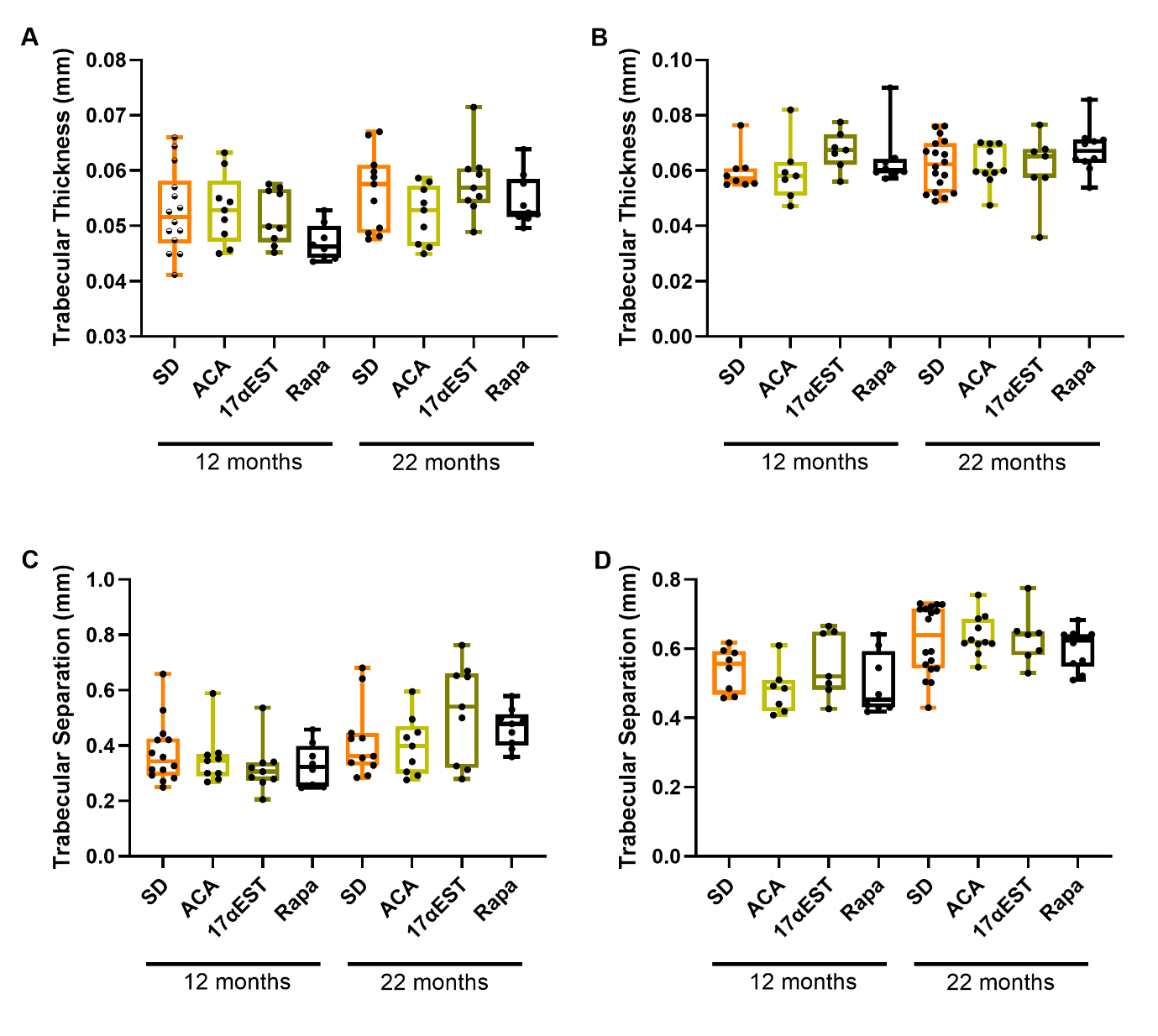

**Supplementary Fig 1.** **Geroprotective interventions do not affect trabecular thickness and separation in UM-HET3** **mice**. Box plot shows the median (center line), interquartile range (box) and maximum and minimum value (whiskers). Each dot represents one animal. A) Trabecular thickness in male UM-HET3 mice receiving standard diet (SD), Acarbose (ACA), 17α-estradiol (17αEST), rapamycin (Rapa), analysed at 12 or 22 months of age. B) Trabecular thickness in female UM-HET3 mice exposed to the same interventions; C) Trabecular separation in the same male mice shown in panel A. D) trabecular separation in the same female mice shown in panel B.

| **Age** | **Intervention** | **Sex** | **Large** | **Medium** | **Small** | **Negligible** |
| --- | --- | --- | --- | --- | --- | --- |
| 12 | 17aEST | Female | 0 | 0 | **0.969** | 0.031 |
| 12 | 17aEST | Male | 0 | 0 | 0 | **1** |
| 12 | ACA | Female | 0 | 0 | 0 | **1** |
| 12 | ACA | Male | 0 | 0 | 0 | **1** |
| 12 | Rapa | Female | 0 | 0 | 0.009 | **0.991** |
| 12 | Rapa | Male | 0 | 0 | **0.926** | 0.074 |
| 22 | 17aEST | Female | 0 | 0 | 0 | **1** |
| 22 | 17aEST | Male | 0 | 0 | 0 | **1** |
| 22 | ACA | Female | 0 | 0 | 0 | **1** |
| 22 | ACA | Male | 0 | 0 | 0.055 | **0.945** |
| 22 | Rapa | Female | 0 | 0 | **0.64** | 0.36 |
| 22 | Rapa | Male | 0 | 0 | 0 | **1** |

**Supplementary table 1**. **Posterior probabilities of standardised effect sizes for trabecular thickness relative to standard diet, classified according to regions of practical equivalence (ROPE) by sex and age.** Rope categories correspond to standardised effects sizes: negligible (absolute value < 0.2), small (0.2–0.5), medium (0.5–0.8), or large (> 0.8). The category with the highest probability in each group is highlighted.

| **Age** | **Intervention** | **Sex** | **Large** | **Med** | **Small** | **Negligible** |
| --- | --- | --- | --- | --- | --- | --- |
| 12 | 17aEST | Female | 0 | 0 | 0 | **1** |
| 12 | 17aEST | Male | 0 | 0 | **0.952** | 0.048 |
| 12 | ACA | Female | 0 | 0 | **0.796** | 0.204 |
| 12 | ACA | Male | 0 | 0 | 0 | **1** |
| 12 | Rapa | Female | 0 | 0 | 0.107 | **0.893** |
| 12 | Rapa | Male | 0 | 0 | **0.569** | 0.431 |
| 22 | 17aEST | Female | 0 | 0 | 0 | **1** |
| 22 | 17aEST | Male | 0 | 0 | **0.95** | 0.05 |
| 22 | ACA | Female | 0 | 0 | 0 | **1** |
| 22 | ACA | Male | 0 | 0 | 0 | **1** |
| 22 | Rapa | Female | 0 | 0 | 0 | **1** |
| 22 | Rapa | Male | 0 | 0 | **0.956** | 0.044 |

**Supplementary figure 2.** **Posterior probabilities of standardised effect sizes for trabecular separation relative to standard diet, classified according to regions of practical equivalence (ROPE) by sex and age.** Rope categories correspond to standardised effects sizes: negligible (absolute value < 0.2), small (0.2–0.5), medium (0.5–0.8), or large (> 0.8). The category with the highest probability in each group is highlighted.
